# Attentive deep learning-based tumor-only somatic mutation classifier achieves high accuracy agnostic of tissue type and capture kit

**DOI:** 10.1101/2021.12.07.471513

**Authors:** R. Tyler McLaughlin, Maansi Asthana, Marc Di Meo, Michele Ceccarelli, Howard J. Jacob, David L. Masica

## Abstract

In precision oncology, reliable identification of tumor-specific DNA mutations requires sequencing tumor DNA and non-tumor DNA (so-called “matched normal”) from the same patient. The normal sample allows researchers to distinguish acquired (somatic) and hereditary (germline) variants. The ability to distinguish somatic and germline variants facilitates estimation of tumor mutation burden (TMB), which is a recently FDA-approved pan-cancer marker for highly successful cancer immunotherapies; in tumor-only variant calling (i.e., without a matched normal), the difficulty in discriminating germline and somatic variants results in inflated and unreliable TMB estimates. We apply machine learning to the task of somatic vs germline classification in tumor-only samples using TabNet, a recently developed attentive deep learning model for tabular data that has achieved state of the art performance in multiple classification tasks (Arik and Pfister 2019). We constructed a training set for supervised classification using features derived from tumor-only variant calling and drawing somatic and germline truth-labels from an independent pipeline incorporating the patient-matched normal samples. Our trained model achieved state-of-the-art performance on two hold-out test datasets: a TCGA dataset including sarcoma, breast adenocarcinoma, and endometrial carcinoma samples (F1-score: 88.3), and a metastatic melanoma dataset, (F1-score 79.8). Concordance between matched-normal and tumor-only TMB improves from *R*^*2*^ = 0.006 to 0.705 with the addition of our classifier. And importantly, this approach generalizes across tumor tissue types and capture kits and has a call rate of 100%. The interpretable feature masks of the attentive deep learning model explain the reasons for misclassified variants. We reproduce the recent finding that tumor-only TMB estimates for Black patients are extremely inflated relative to that of White patients due to the racial biases of germline databases. We show that our machine learning approach appreciably reduces this racial bias in tumor-only variant-calling.

## Introduction

An important application of somatic variant calling is patient selection in cancer immunotherapy clinical trials because somatic mutation count can predict response to immune checkpoint inhibitors (ICI)(Samstein et al. 2019; Wu et al. 2019; Litchfield et al. 2021). Tumor mutation burden (TMB)—defined as the number of coding nonsynonymous somatic mutations per megabase of DNA, and often measured through whole exome sequencing (WES)—is a strong predictor of response and survival in solid tumors. Encouraged by the recent results from the successful phase 2 KEYNOTE-158 trial(Marabelle et al. 2020), the FDA has approved TMB as a marker across all tumor subtypes for the anti-PD1 ICI pembrolizumab, where increased TMB is associated with increased benefit. This approval broadens the importance of reliably estimating patient-level TMB using WES data.

In addition to TMB, identifying somatic and germline variants is used to understand the molecular basis of cancer. Somatic mutation underlies cancer formation and progression, often through gain-of-function mutations in oncogenes and loss-of-function mutations in tumor suppressors(Martincorena and Campbell 2015). It is becoming increasingly crucial to characterize and identify somatic mutations to predict whether a cancer patient will be resistant or responsive to existing targeted therapies. Germline variation in genes such as BRCA and TP53 can also be heritable cancer drivers, so understanding the germline context of cancer can complement the characterization of acquired somatic mutations.

Matched-normal samples are not always available in the clinic, leading to entirely tumor-only cohorts and mixed cohorts of tumor-only and matched-normal samples. Causes for missing a matched normal sample include failed quality control in the normal samples and a lack of consent to procure germline blood samples. Furthermore, acquisition of a patient’s matched normal must be included in the design of the oncology clinical trial, which is not a routine practice.

The absence of a patient-matched normal complicates somatic variant calling in precision oncology. The sheer number of rare germline variants per sample and their broad distribution of variant allele fractions (VAFs) (Supplemental Figure 1) makes it a challenge to retrieve the relatively small number of genuine somatic mutations. One study reported the absence of a matched normal sample leads to a 67% false positive rate; thus, most putative somatic mutations in tumor-only variant calling are actually rare germline variants(Shi et al. 2018). The resulting tumor-only TMB estimate is artificially inflated relative to that derived via germline variant subtraction using a matched normal. One recent study reported a fold inflation of 2.2-16.9 for tumor-only-calculated TMB, depending on the chosen germline database filtering strategy(Parikh et al. 2020).

Several computational methods have been developed to improve tumor-only variant calling, either by sophisticated filtering approaches(Sukhai et al. 2019), or via explicit statistical inference of the somatic alteration state of the cancer genome (as in the algorithms ABSOLUTE(Carter et al. 2012) and CLONET(Prandi et al. 2014)). The latter category include PureCN(Riester et al. 2016; Oh et al. 2020) and SGZ(Sun et al. 2018), two recently developed Bayesian methods that infer the altered genomic state of the tumor to estimate somatic and germline probabilities in samples without a matched normal. These methods first estimate global properties of the cancer genome--purity and ploidy--as well as local DNA copy number. They integrate this information with the observed variant allele frequencies (VAFs) to calculate the posterior probability that a mutation is somatic. The complexity of the cancer genome, including clonality and structural variation, coupled with the complex statistics of next-generation sequencing(Poplin et al. 2018) makes improving upon these statistical models challenging. Recently, state-of-the-art speed and accuracy have been achieved using machine learning for somatic variant calling with matched-normal samples(Wood et al. 2018; Huang et al. 2019; Sahraeian et al. 2019). Rather than attempting to model explicitly the likelihood functions for somatic mutations, these methods involve training a machine learning classifier on a diverse training set with truth labels and applying the trained classifier to new oncology samples. Taking inspiration from these studies, we hypothesized a supervised machine learning algorithm would be effective for classifying mutations as somatic or germline in patient derived solid tumor samples lacking a matched normal.

## Results

### Train / test overview

We used a somatic mutation calling pipeline to process samples both with and without the matched normal sample (see Materials and Methods). Because TMB, copy-number variation (CNV), and sample composition (tumor purity) can impact somatic mutation calling, we selected oncology samples from different tissue types that span biological extremes, including: ovarian adenocarcinoma (high purity, low TMB, high CNV), sarcoma (low TMB, high CNV), testicular germ cell cancer (extremely low TMB), endometrial carcinoma, colorectal adenocarcinoma, metastatic melanoma, and lung adenocarcinoma and squamous carcinoma (high TMB), and several other cancer subtypes from the Cancer Genome Atlas (TCGA).

### Model

We engineered 30 mutation- and copy-number-specific features using tumor-only samples (see Materials and Methods). This included traditional features for somatic variant calling such as germline database frequency, COSMIC somatic mutation database counts, and read-based statistics such as variant allele fraction (VAF) and major allele frequency. Expecting somatic mutations to exhibit a different mutational spectrum from germline variants, we also included features that characterize the trinucleotide context and base substitution subtypes that are the basis for mutational signature analysis(Alexandrov et al. 2020). The local copy number for each variant is represented by features derived from copy-number segmentation data and variant calls. Briefly, using germline variant databases and copy-number segments, we identify neighboring heterozygous germline SNPs of similar copy number, and bin the variant counts into 20 non-overlapping VAF bins (see Materials and Methods).

The somatic and germline truth labels were determined by running an independent variant-calling pipeline using the matched normal samples. Variants passing in the matched-normal pipeline were considered somatic; all other variants in the tumor-only pipeline were considered germline. The merged tumor-only feature matrix and truth labels were used for the binary somatic vs germline classification task.

To classify mutations, we selected TabNet, an attentive deep learning model for tabular data(Arik and Pfister 2019) to classify mutations as somatic or germline. TabNet leverages attention modules and has been shown to achieve state-of-the-art performance on tabular data, outperforming XGBoost and other powerful supervised machine learning models. In addition to improved accuracy over XGBoost, the feature masks of TabNet allow interpretation for each classification instance and allow the saliency of each feature and each instance to be visualized in a matrix. The overall train / validation / test scheme is illustrated in Figure 1.

**Figure 1.**
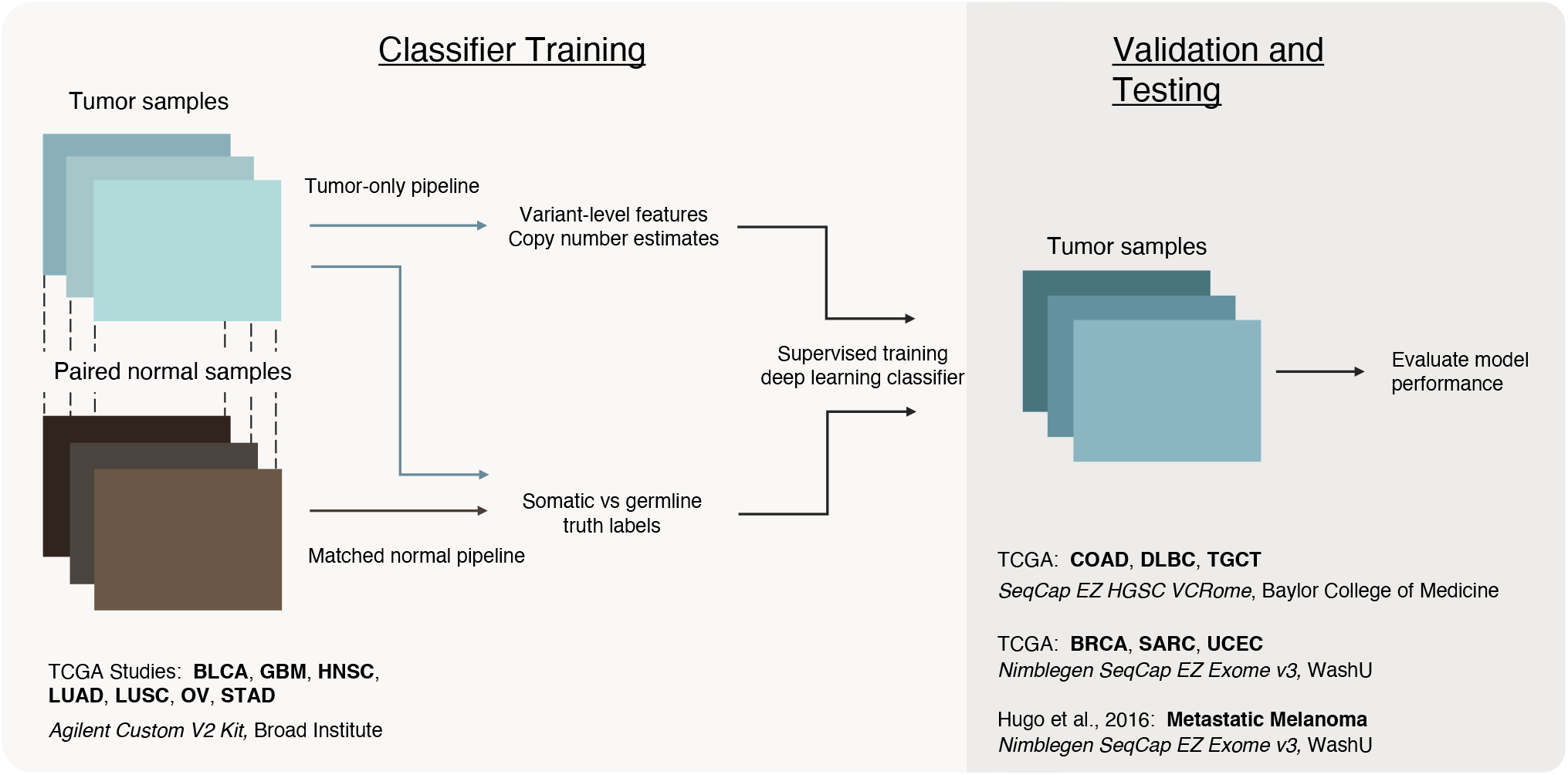
Methodology: somatic vs germline prediction using supervised learning. To improve the reliability of tumor-only variant calling in whole exome sequencing (WES) samples, we classify mutations as ‘somatic’ or ‘germline’ using TabNet, an attentive deep learning classifier for tabular data. We train TabNet on solid tumor data in a supervised manner, using features from a tumor-only analysis and truth-labels derived from a matched-normal analysis. We evaluate the trained model blindly on hold-out tumor sample data that is biologically and technically distinct from the training set data– i.e., from different tissues of origin, exome capture-kits, and sequencing centers. **Classifier Training** *(left)*: To prepare the training set, we first align WES data for tumor and matched normal samples from 105 oncology patients in The Cancer Genome Atlas across 7 studies (BLCA = Bladder Urothelial Carcinoma, GBM = Glioblastoma Multiforme, HNSC = Head and Neck Squamous Cell Carcinoma, LUAD = Lung Adenocarcinoma, LUSC = Lung Squamous Cell Carcinoma, OV = Ovarian Serous Cystadenocarcinoma, STAD = Stomach Adenocarcinoma). For all patients, variant calling is performed with and without the matched normal reference. CNV analysis is performed without the matched normal samples. We extract features from the tumor-only variants and CNV data. The somatic or germline status of each variant detected in the matched normal variant calling pipeline is used as the ground truth label--0 for germline and 1 for somatic. We combine the features and truth labels to train a TabNet classifier to distinguish somatic from germline variants. **Validation and Testing** *(right):* The model with the best recall and precision on a validation set (COAD, DLBC, and TGCT, SeqCap EZ HGSC VCRome, Baylor College of Medicine) is selected and applied to two hold-out test datasets of tumor-only samples: TCGA BRCA, SARC, and UCEC, and the metastatic melanoma data set of Hugo et al., (2016), using the Nimblegen SeqCap EZ Exome v3 kit. Accuracy and run-time of the tumor-only classification method are benchmarked using truth labels from the associated matched-normal pipeline.

### Training set construction

For our training set we selected 105 tumor samples from distinct patients in seven cancer subtypes from TCGA. Somatic and germline truth labels were generated using the results of a variant-calling pipeline that included the patient-matched normal samples. We engineered features for our deep learning classifier using the variant and CNV calls from the independent tumor-only pipeline, which used a process-matched panel of normals for each patient that did not include the patient-matched normal samples (see Materials and Methods). The training dataset consisted of 15 samples from each of seven solid tumor cancer subtype studies in the Cancer Genome Atlas: bladder urothelial carcinoma (BLCA)(Cancer Genome Atlas Research Network 2014a), glioblastoma multiforme (GBM)(Brennan et al. 2013), head and neck squamous cell Carcinoma (HNSC)(Cancer Genome Atlas Network 2015), lung adenocarcinoma (LUAD)(Cancer Genome Atlas Research Network 2014b), lung squamous cell carcinoma (LUSC)(Cancer Genome Atlas Research Network 2012), ovarian serous cystadenocarcinoma(OV)(Cancer Genome Atlas Research Network 2011), and stomach adenocarcinoma (STAD)(Cancer Genome Atlas Research Network 2014c). To impose technical consistency within the training set, all the selected samples from these studies were sequenced at the Broad Institute using the Agilent Custom V2 exome capture kit. This was done so we could later investigate whether our trained model could generalize to other exome capture kits and still achieve high classification accuracy.

### Validation set construction

Our validation set consisted of 15 tumor samples from each of three cancer subtypes (45 total): colon adenocarcinoma (COAD)(Cancer Genome Atlas Network 2012a), lymphoid neoplasm diffuse large B-cell lymphoma (DLBC), and testicular germ cell tumors (TGCT)(Shen et al. 2018). These three TCGA cohorts were sequenced at Baylor College of Medicine with the SeqCap EZ HGSC VCRome capture kit. These validation WES data samples have been shown to exhibit different biases in genomic coverage than the Agilent Custom V2 kit(Wang, Kim, and Chuang 2018). TabNet classified the variants in the full validation set. The resulting predictions were subsequently compared to the matched-normal truth labels and evaluated for performance (Table 1; Figure 2). After 100 training epochs, the optimal model was selected as the model with the best classification performance on the validation data. Thus, we selected the model with the best generalization to new tumor types sequenced with a new capture kit.

**Table 1).** Benchmark accuracy metrics for somatic vs germline classification by TabNet and PureCN. Overall performance considers all single-nucleotide variants (SNVs) and indels. TP: true positives – somatic mutations correctly classified as somatic. FP: false positives; rare germline variants misclassified as somatic mutations. FN: false negatives – somatic mutations misclassified as germline variants. TN: true negatives – rare germline mutations correctly classified as germline. Call rate – percentage of total coding variants classified.

| Method | Dataset | Variant category | AUC | Sensitivity | Specificity | Positive predictive value | Negative predictive value | F1-score | Balanced accuracy | FP | TP | FN | TN | Total classified variants | Call rate |
| --- | --- | --- | --- | --- | --- | --- | --- | --- | --- | --- | --- | --- | --- | --- | --- |
| TabNet | Blind test set - BRCA SARC UCEC | Overall | **0.942** | **0.931** | 0.828 | 0.839 | **0.926** | **0.883** | **0.879** | 5397 | 28184 | 2086 | 25962 | 61629 | 100 |
|  |  | SNVs | **0.945** | **0.943** | 0.827 | 0.843 | **0.937** | **0.89** | **0.885** | 5199 | 27918 | 1675 | 24923 | 59715 | 100 |
|  |  | Indels | 0.784 | **0.838** | 0.624 | 0.515 | **0.89** | 0.638 | 0.731 | 662 | 703 | 136 | 1100 | 2601 | 100 |
|  | Blind test set - metastatic melanoma | Overall | **0.852** | **0.797** | **0.753** | **0.799** | **0.751** | **0.798** | **0.775** | 3176 | 12598 | 3215 | 9695 | 28684 | 100 |
|  |  | SNVs | **0.85** | **0.796** | **0.749** | **0.8** | **0.744** | **0.798** | **0.773** | 3114 | 12490 | 3198 | 9298 | 28100 | 100 |
|  |  | Indels | 0.82 | 0.837 | 0.619 | 0.346 | 0.94 | 0.489 | 0.728 | 233 | 123 | 24 | 378 | 758 | 100 |
| PureCN | Blind test set - BRCA SARC UCEC | Overall | 0.85 | 0.662 | **0.912** | **0.882** | 0.729 | 0.756 | 0.787 | 2237 | 16792 | 8588 | 23065 | 50682 | 82.2 |
|  |  | SNVs | 0.851 | 0.662 | **0.913** | **0.887** | 0.724 | 0.758 | 0.788 | 2082 | 16343 | 8360 | 21983 | 48768 | 81.7 |
|  |  | Indels | **0.827** | 0.606 | **0.893** | **0.756** | 0.805 | **0.673** | **0.749** | 132 | 410 | 267 | 1105 | 1914 | 73.6 |
|  | Blind test set - metastatic melanoma | Overall | 0.824 | 0.789 | 0.732 | 0.794 | 0.726 | 0.791 | 0.76 | 2951 | 11340 | 3041 | 8050 | 25382 | 88.5 |
|  |  | SNVs | 0.823 | 0.788 | 0.729 | 0.797 | 0.717 | 0.792 | 0.758 | 2856 | 11228 | 3028 | 7686 | 24798 | 88.2 |
|  |  | Indels | **0.887** | **0.888** | **0.826** | **0.581** | **0.964** | **0.703** | **0.857** | 80 | 111 | 14 | 379 | 584 | 77 |

**Figure 2.**
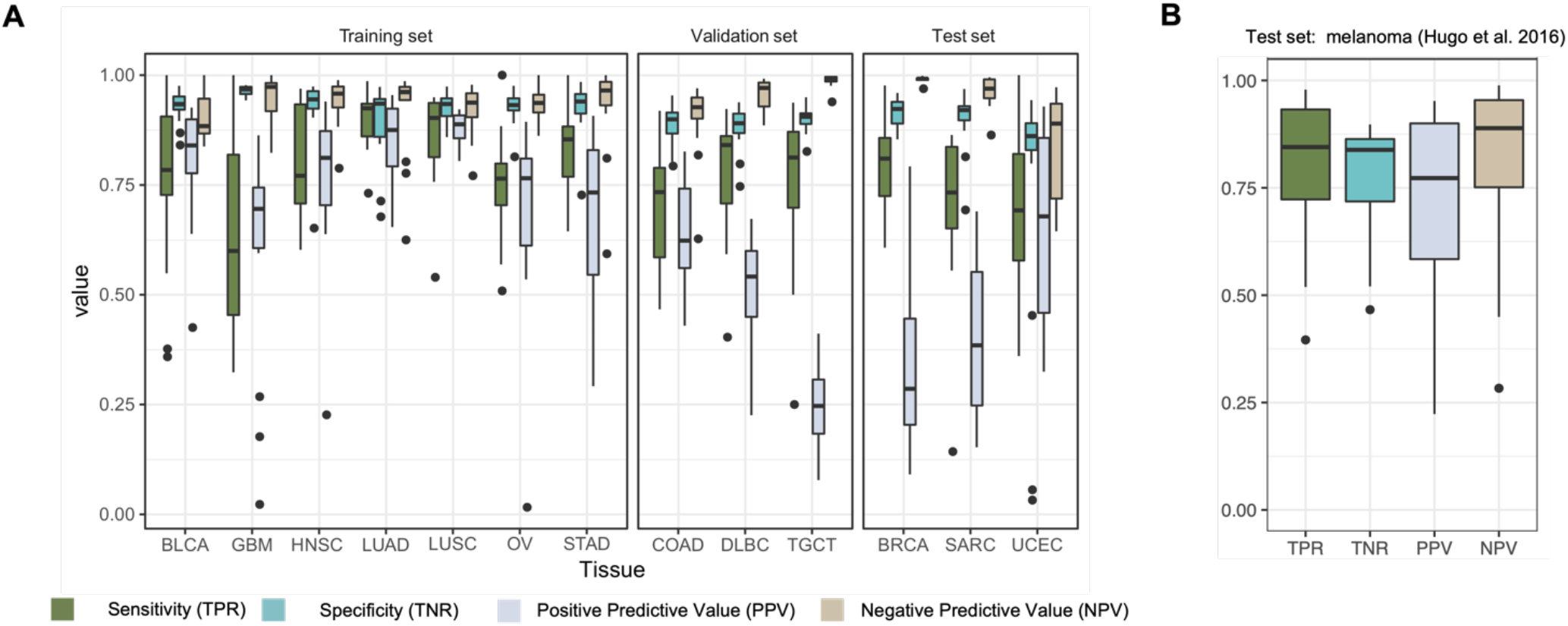
Classifier’s patient-level performance stratified by tissue type. **A)** Four accuracy metrics for TCGA datasets. n = 15 randomly selected cancer patients per tissue type. TabNet deep learning classifier trained on BLCA, GBM, HNSC, LUAD, LUSC, OV, STAD studies, all with Agilent Custom V2 exome capture kit *(left)*, validated on COAD, DLBC, and SARC studies, with SeqCap EZ HGSC VCRome capture kit (*center*, n = 45), and tested on BRCA, SARC, and UCEC Nimblegen SeqCap EZ v3 capture kit (*right*, n = 45). **B)** Four accuracy metrics for the Hugo metastatic melanoma data set (23 cancer patients sequenced by UCLA).

### Training and validation results

During training, the optimal trained model was selected as the one with the best performance on the validation set. This optimal model achieved an AUC of 0.962 on the training data. After choosing the best posterior probability threshold for somatic vs germline classification, the model achieved an F1-score of 0.859, sensitivity (true positive rate, TPR) of 0.869 and positive predictive value (PPV) of 0.848. These results suggest the model was fitting the data yet not overparameterized to the point of memorizing the training data. The performance on the validation data was lower than the training set, with an AUC of 0.909 and an F1-score of 0.633 (TPR: 0.734, PPV: 0.557), suggesting either the model is slightly over-fitting, or the validation data are more challenging to classify. This performance is still a marked improvement over the naive tumor-only method (using a process matched panel of normals and multiple germline databases to remove likely germline SNPs) where all variants remaining after filtering are classified as somatic: F1-score of 0.28 (TPR 1.0, PPV: 0.16). Figure 2A displays patient-level performance for all TCGA datasets, grouped by tissue type. During training, TabNet fit LUAD and LUSC tissues most easily, with the highest TPR and PPV, meaning somatic and germline variants should be relatively easy to discriminate in these cancer types. GBM was the hardest solid tumor subtype to classify, with the lowest TPR and PPV. Thus, tumor-only variant calling appears to be relatively tractable for lung cancers and challenging for GBM. In the validation set, TGCT exhibited high sensitivity and the lowest PPV. COAD exhibited the highest PPV and the lowest sensitivity. The reason for these differences is discussed below in “Explaining variability in performance”.

### Holdout test sets

After model training and selection, we constructed two separate holdout test sets, including four cancer subtypes and a new exome capture kit, Roche Nimblegen SeqCap EZ Exome v3. Results are shown in Table 1.

The first holdout test set included solid tumor samples from 45 patients from the following three TCGA studies (15 each): breast invasive carcinoma (BRCA)(Cancer Genome Atlas Network 2012b), sarcoma (SARC)(Cancer Genome Atlas Research Network 2017), and uterine corpus endometrial carcinoma (UCEC)(Cancer Genome Atlas Research Network 2013). These samples were sequenced at Washington University in St. Louis. Table 1 displays the trained model’s performance on the hold-out test datasets. TabNet achieved an AUC of 0.942, F1-score of 0.883, and a balanced accuracy of 0.879 overall, including SNVs and indels. TabNet performs better on SNVs (AUC: 0.945, F1: 0.89) than indels (AUC: 0.784, F1: 0.638). Across the three tissue subtypes, the best PPV and worst TPR were both observed for UCEC, and the best TPR and worst PPV were both observed for BRCA.

To further identify any potential batch effects, we acquired a final holdout dataset comprising non-TCGA data. This final holdout dataset included 23 samples from the Hugo et al., 2016 metastatic melanoma study(Hugo et al. 2016). Relative to the TCGA test set, our model performed better in both TPR and PPV on this dataset at the patient level (Figure 2B). Yet performance on the overall variant-level was slightly worse due to the influence of high-TMB patients. AUC was 0.852, 0.85, 0.82, F1-score was 0.798, 0.798, and 0.728 for overall, SNVs, and indels, respectively.

### Concordance of tumor-mutational burden estimation methods

The reliable estimation of TMB is a critical benchmark for a model designed to improve tumor-only variant calling. This capacity is especially relevant in immuno-oncology clinical trials where TMB is a potent biomarker of response(Goodman et al. 2017) and survival(Samstein et al. 2019). We define the naive tumor-only method as a non-machine-learning approach that incorporates a process-matched panel of normals, multiple germline variant databases, and standard variant filtering techniques (see Materials and Methods) to remove germline variants and artifacts. Our machine-learning-based approach applies TabNet’s somatic vs germline classifications to the results of the naive approach, using the optimal somatic posterior probability cutoff of 0.5 to isolate predicted somatic mutations.

To evaluate clinical utility, we compared the naïve and machine-learning-based TMB estimates with those derived from the matched-normal gold standard (Figure 3). Using linear regression, we calculated an *R*^*2*^ of 0.156, 0.318, and 0.006 for TCGA train, validation, and test sets, respectively, indicating a weak correlation between matched and tumor-only TMBs (Figure 3A). Notably, in the BRCA, SARC, and UCEC test set, the rank order of TMBs is markedly different between naive and matched-normal methods. The slopes of these fits (0.148, 0.254, 0.016 for TCGA training, validation, and test datasets, respectively) are substantially less than 1.0 in all cohorts, indicating a consistently inflated TMB result for tumor-only samples with a magnitude that agrees with recently reported results(Parikh et al. 2020; Shi et al. 2018). Next, we evaluate the relationship between TMB from TabNet-predicted somatic predictions to matched-normal germline-subtracted TMB (Figure 3B). Linear regression fits for train, validation, and test sets yielded *R*^*2*^ values of 0.938, 0.871, 0.705, indicating a 3-to 44-fold improvement over the naive method. The slope of best fit was similarly encouraging (train, 0.97; validation, 0.913; test, 0.804), with our model achieving a 5-fold improvement relative to the naïve approach. TabNet’s improvement to these concordance metrics argues for the use of TabNet-corrected TMB estimates for clinical variant analysis in tumor-only samples. Importantly, TabNet enables reliable TMB comparisons in mixed cohorts composed of both tumor-only and paired tumor-normal specimens.

**Figure 3.**
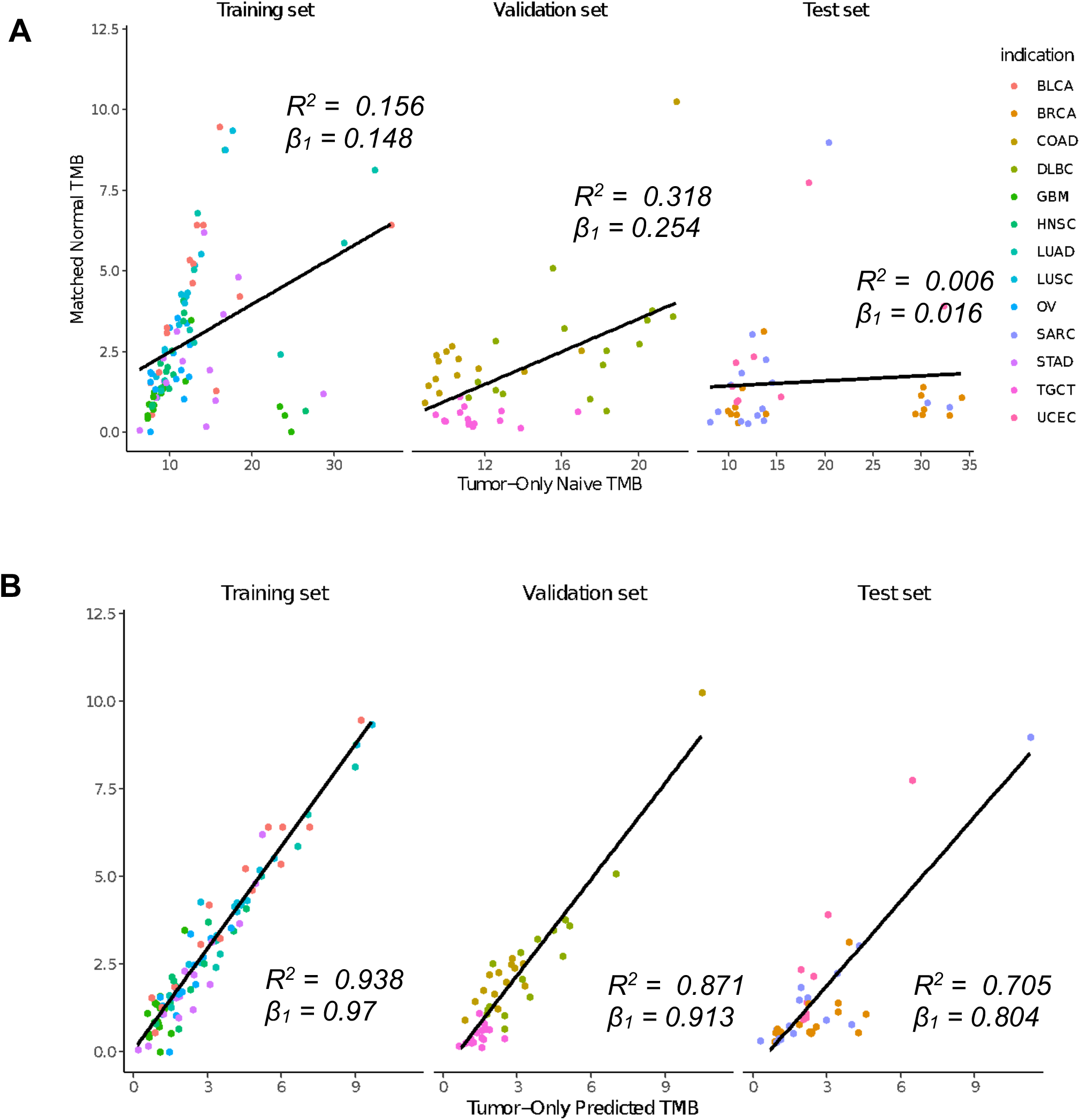
Concordance of Tumor Mutational Burden (TMB) calculated with and without matched normals. Training set, n = 105; Validation set, n = 45; Test set, n = 45 patients. **A)** Matched-normal TMB compared to TMB calculated by naïve tumor-only approach – variants are filtered by removing common germline variants using multiple population germline databases and a process-matched leave-one-out panel of normals. **B)** Matched-normal TMB compared to TMB calculated with TabNet-predicted somatic mutations. *β*_*1*_ indicates the slope of linear regression fit.

### Comparison to PureCN

By varying the posterior probability threshold across 500 quantiles, we constructed ROC and precision-recall curves for TabNet and PureCN. On the training data, we found optimal F1-scores of 0.868 near a threshold of 0.48 for TabNet, and 0.842 near a threshold of 0.0051 for PureCN. However, after breaking down performance into SNVs and indels, we noticed different probability thresholds yielded optimal results for the two variant categories, for both TabNet and PureCN. For SNVs, TabNet’s optimal F1-score occurred at a cutoff of 0.508, PureCN’s at 0.005. For indels, the optimal F1-score occurred at a threshold of 0.1368 for Tabnet and 0.014 for PureCN. We used these probability thresholds derived from performance on the training set to make our binary predictions for the blind test sets. These are the results reported in Table 1. Figure 4A displays the ROC curve comparing TabNet and PureCN for the BRCA, SARC, and UCEC TCGA holdout test set of 45 patient samples. Both algorithms are highly tunable and TabNet’s has higher AUC and is consistently concave down, suggesting more stable dependence on posterior-probability cutoffs.

**Figure 4:**
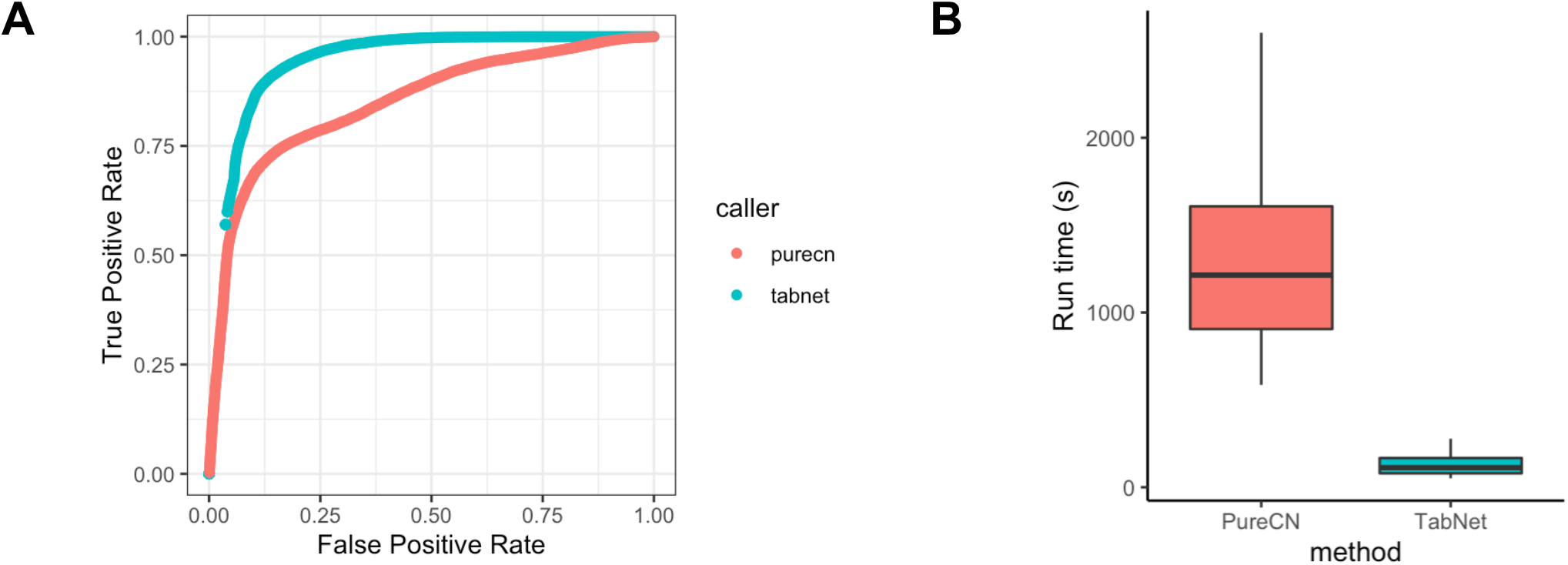
Comparison between PureCN and TabNet on TCGA holdout test set. **A)** Receiver operating characteristic (ROC) curve calculated for both TabNet and PureCN, treating somatic mutations as positives and germline variants as negatives. Curves display 500 distinct posterior probability thresholds for classification, selected by binning the probabilities into 500 quantiles. **B)** Run-time comparison in seconds. (PureCN, 250 CPUs per sample), TabNet 1 CPU per sample, no GPU)

For the holdout datasets, TabNet achieved better overall performance and better performance on SNVs, with PureCN achieving better performance on indels. On the TCGA holdout test set, TabNet has the best F1-score overall and for SNVs, with a 12.7% improvement overall and 13.2% improvement for SNVs relative to PureCN. For indels, PureCN achieved an F1-score of 67.3% and 74.9% balanced accuracy, whereas Tabnet achieved an F1-score of 63.8% and 73.1% balanced accuracy. On the second hold-out test set consisting of 23 metastatic melanoma patients, TabNet performs nearly identically to PureCN, both overall (better by 0.7%) and for SNVs (better by 0.6%) but PureCN’s performance is substantially better on indels by 21.4%, with 153 fewer false positives than TabNet and only 12 fewer true positives.

Next, we compared the amount of time elapsed to make somatic/germline predictions starting from annotated tumor-only VCFs. Compute time of TabNet (mean 133.7 seconds) on a single core was 9.6 times faster than PureCN’s (1287.3 seconds, *p* << 0.001) using 250 cores (Figure 4B). This dramatic speed improvement over PureCN is not surprising as Bayesian methods are known to require intense CPU resources.

### Global feature importance

We inspected the global feature importances of our trained classifier. The top 30 out of 56 total features are shown in Figure 5A. The maximum population allele frequency across multiple germline databases (*pop_max*) is the most important feature, with an importance score of 0.49. The next most important features are *t_maj_allele* (the greatest VAF among all observed alleles at the variant’s locus, 0.13), *max_cosmic_count (*number of times the variant is observed in COSMIC, the catalog of somatic mutations in cancer(Tate et al. 2019), 0.12), *t_alt_freq* (the VAF of the mutant allele, 0.09), snp_vaf_bin_00 (the number of neighboring heterozygous SNPs with VAF between 0.0 and 0.05, 0.022), and *count* (the total number of variants to classify in the sample containing the variant, 0.019). All features are variant-specific except *count*, which has the same value across all variants in each patient. *inframe_indel* (importance = 0.0090) is the only ontology-related feature in the top 30. The remaining features in the top 30 are from either the *snp_vaf_bin* class of features or are related to the mutational spectrum. The *snp_vaf_bin* features (derived from binning the VAFs of neighboring heterozygous common SNPs) together add up to 0.067. The features characterizing the mutational spectrum--substitutions such as *C>G* and trinucleotide contexts like *TTG* -- together add up to a feature importance of 0.066.

**Figure 5:**
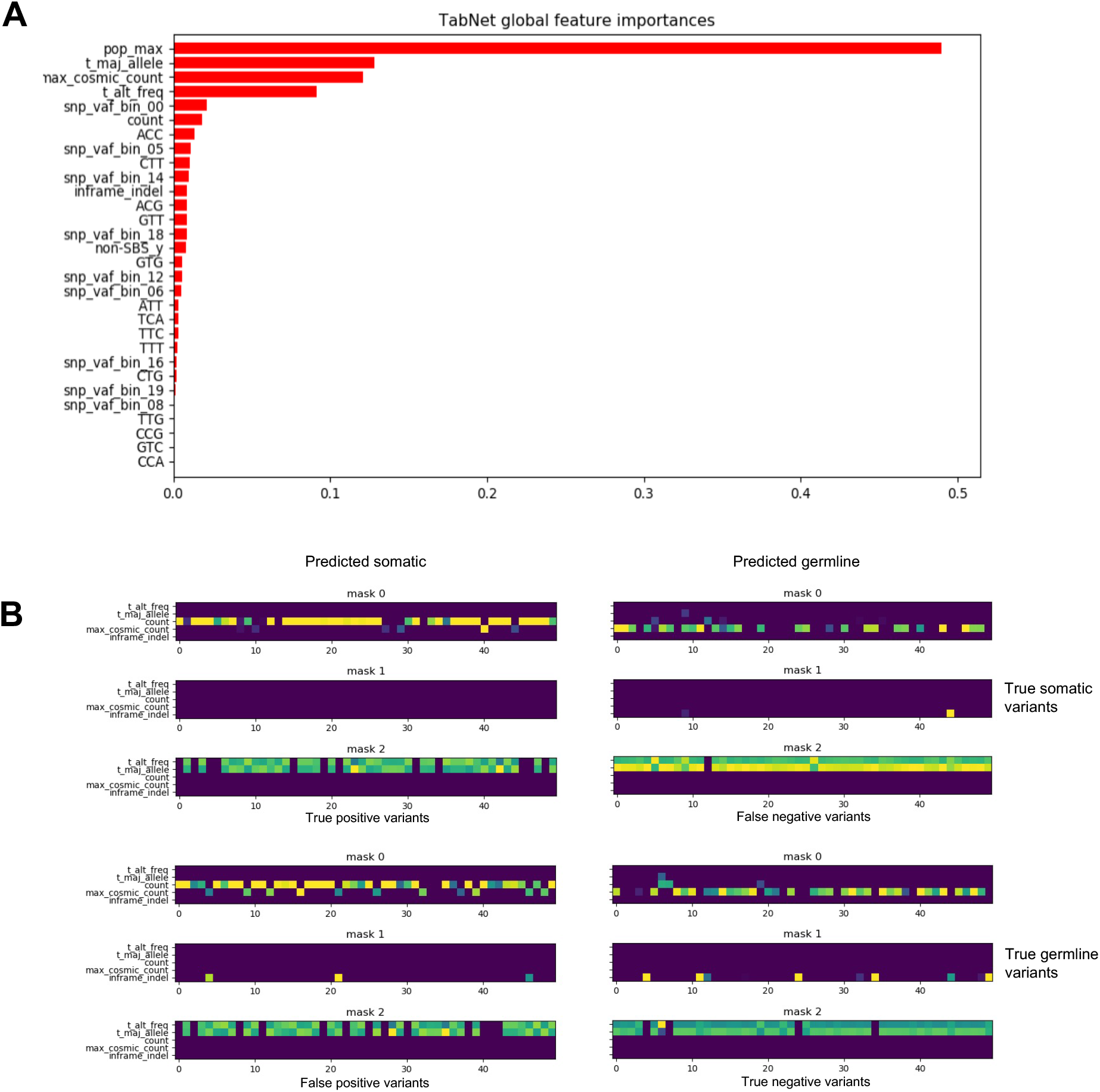
Global and local feature importance for trained TabNet somatic vs germline classifier. **A)** ranked global feature importance. *pop_max*: maximum population frequency of variant across multiple germline databases, *t_maj_allele*: the fraction of reads containing the most supported allele at that locus in the sample, *max_cosmic_count*: the number of occurrences of variant in COSMIC somatic database, *t_alt_freq*: the fraction of reads supporting the alternate allele, *snp_vaf_bin_i*: the number of informative SNPs in the copy-number segment with a VAF between i/20 and (i+1)/20, *count*: the total number of variants to classify in that sample,, *ACC, CTT, etc*: trinucleotide context, *non-SBS-y*: non single-base substitution so no trinucleotide context applies. **B) TabNet’s f**eature masks explain how the neural network allocates its attention during classification. For each of the four categories – true positives (*left*), false positives (*top right*), false negatives (*bottom left*), true negatives *(bottom right) –* 50 variants are randomly selected from the TCGA hold out test set. Only the first 3 feature masks are shown, and only the first 5 variables are shown.

Together, these features are reasonably ordered, with *pop_max* being the most important, and allele fraction and COSMIC features near the top. A surprise is the low feature importance of the *nonsense* ontology. This is a surprise because nonsense mutations are expected to occur more commonly as somatic mutations than as germline variants. Indeed, 63% of the nonsense variants in our test set are truly somatic whereas for all variants in our test set, the truth labels are 49% somatic and 51% germline. The ontology features including *missense, nonsense*, and *inframe_indel* add up only to 0.009. But otherwise, the global feature importances for TabNet appear to be well-ordered.

### Explaining variability in performance

Using multiple regression models, we explain individual performance as a function of several predictor variables. We observed that the most influential factor on a sample’s positive predictive value (PPV) is the “true” TMB coming from germline subtraction via the matched-normal pipeline. Samples with lower TMB tend to have a lower PPV (*R*^*2*^ = 0.54, Supplemental Figure 2), so a paucity of true somatic mutations in a sample contributes to a low PPV. We suspected that a small proportion of germline variants appear somatic based on low variant allele fraction (VAF) leading to a constant level of false positives. Supporting this, the median VAF of false positive (FP) and true negative variants (TN) was 0.35, and 0.50, respectively.

The interpretable feature masks of the TabNet architecture allow us to explain explicitly which features contribute most to these false positive calls. By examining the feature masks, which indicate where the TabNet neural network is allocating its attention for each classification instance, we see surprisingly that the VAF (*t_alt_freq*) is not the most distinguishing feature between FPs and TNs. Rather, it is the COSMIC count (*max_cosmic_count*, the number of times the variant is observed in COSMIC) and the overall count of mutations (*count* variable) that best distinguish somatic and germline predictions, with differences being present in the first feature mask layer (Figure 5B). Illustrating the explanatory utility of these feature masks, we found the proportion of variants with a nonzero max_cosmic_count was significantly greater for FPs than for TNs (*p* < 0.00001, Fisher’s exact test), with 993 out of 5397 for FPs (18%) and 3612 out of 25962 (14%) for TNs, and further, the mean max_cosmic_count values was lower for FPs (0.99) than for TNs (0.26), *p* << 0.001 (Wilcoxon rank sums test). We also found that the number of mutations to classify (*count* variable, rare germline SNPs + true somatic mutations in the sample) was greater in FPs (3390) than for TNs (2287), p << 0.001. Thus, significant differences are found between correctly and incorrectly classified germline variants for both features. Together this exemplifies how the feature masks of TabNet help with interpreting classifications.

Considering FPs, TNs, alongside true positives and false negatives, as shown in Figure 5B, the attention of the TabNet classifier is consistently applied to *max_cosmic_count* for all mutations classified as somatic (TP + FP), and to *count* for all variants classified as germline (TN + FN). Limited interpretability is a common and valid criticism of non-attention-based deep learning models, but attention-derived insights such as those presented here and by others(Chen et al. 2020) offer a way to interrogate deep learning models and avoid reliance on predictions from “black box” neural networks.

The sensitivity (TPR) of our model is best explained by the median VAF of the true somatic mutations (MVTSM), a metric that can be interpreted as an approximation for tumor purity divided by 2 (Supplemental Figure 3). The regression model for TPR with MVTSM as the sole predictor has the following parameters: *R*^*2*^ = 0.4, *β*_*0*_(*y*-intercept) = 0.99, *β*_*1*_(slope) = -0.89. Thus, the following approximation predicts the TPR of our model:

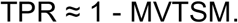

Another interpretation is that the TPR increases with increasing stromal fraction, the so-called “contaminating normal tissue” in the biopsy. Covariance analysis of TPR vs MVTSM across indications identified GBM, melanoma, and BLCA as the tissue types in our datasets where this relationship is strongest (Supplemental Figure 4).

### Impact of racially biased germline databases in tumor-only variant calling

The presence of racial biases in genomic databases has widespread negative implications in human genome science and has been the subject of intense criticism(Popejoy and Fullerton 2016; Bentley, Callier, and Rotimi 2020). Tumor-only variant calling is no exception(Halperin et al. 2017). A recent study observed that the inflation of TMB caused by the absence of a matched normal sample is most severe in underrepresented minorities(Asmann et al. 2021). Comparing the “true” matched-normal TMBs of the 12 Black patients and 55 White patients in the TCGA validation set and hold-out test set, we see no statistical difference in TMB between the two groups (*p* > 0.05, Wilcoxon test) (Figure 6A). In the absence of a matched normal sample, however, the difference is profound (*p* << 0.001) with median tumor-only TMBs of Blacks (30.36) being almost three times as high as that for Whites (11.15) (Figure 6B). After applying TabNet’s somatic germline classifications, the corrected median TMBs for Blacks and Whites are 3.43 and 1.85, respectively. The TabNet-corrected tumor-only TMB calculation for Blacks is still somewhat over-inflated, but the TMB difference of ∼1.6 between Black and White patients constitutes a dramatic reduction of bias relative to the TMB difference of ∼19 seen with the naive tumor-only method (Figure 6C).

**Figure 6:**
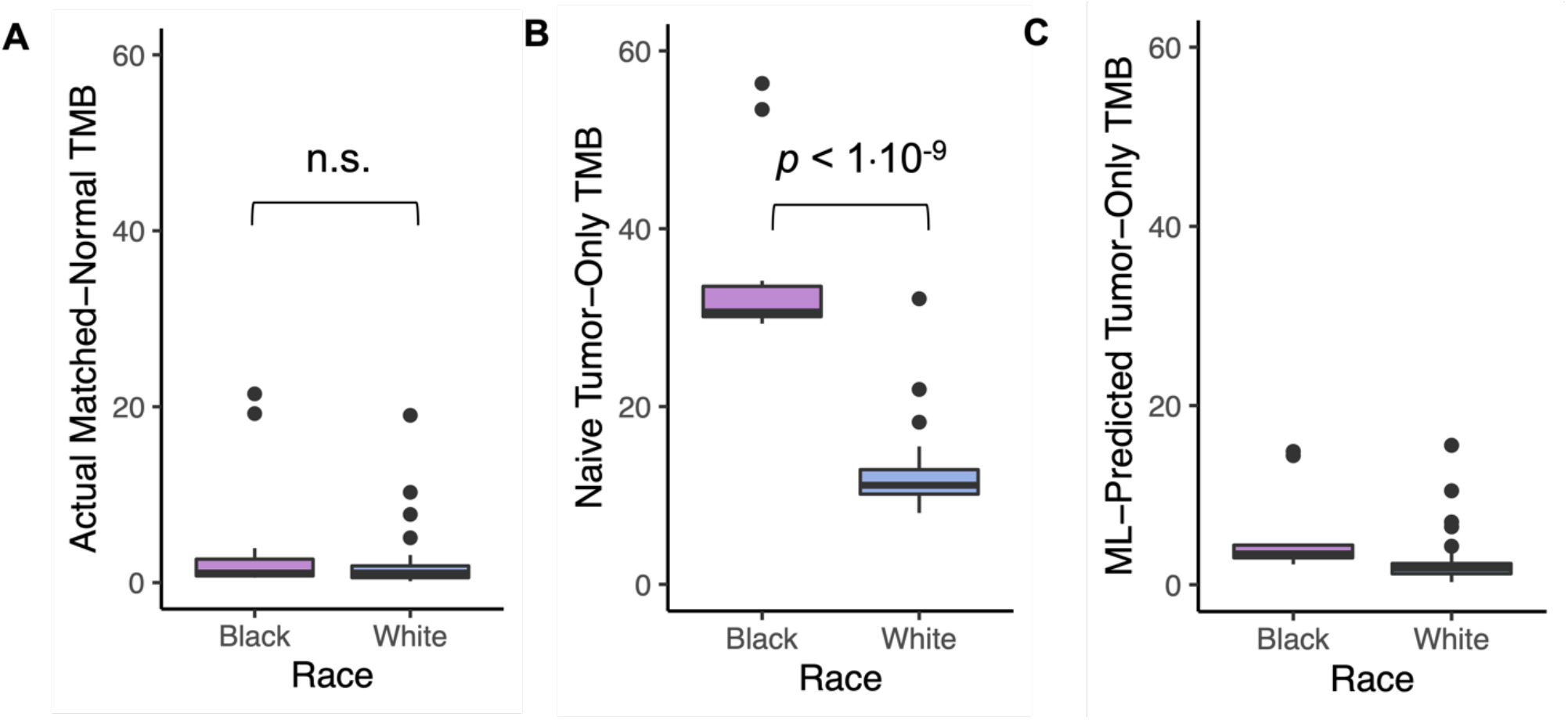
Impact of racial bias in germline databases on Tumor Mutational Burden (TMB) estimates in tumor-only WES samples. Each panel displays patients from TCGA validation and hold-out test sets, n = 12 Black and 55 White patients. **A)** True TMB from matched normal pipeline. **B)** TMBs without matched normal samples, using multiple germline population databases and a process-matched leave-one-out panel of normals (naïve tumor-only method). B) TabNet-corrected tumor-only TMBs.

## Discussion

We constructed and trained an attentive deep learning classifier to distinguish somatic mutations from rare germline variants. Our model was trained on seven cancer subtypes sequenced by the Broad Institute with a single exome capture kit. This trained model generalizes to two distinct capture kits and seven distinct cancer subtypes in the validation and two blind hold-out sets. TabNet outperforms PureCN in both speed and accuracy. TabNet predictions confer agreement between matched and tumor-only TMB calculation, with a fold improvement as much as 44 over the naive method and a slope within 20% of 1.0, enabling reliable somatic mutation retrieval in tumor-only variant calling and harmonization of TMB calculation in cohorts of mixed tumor-only and matched-normal WES samples.

The performance metrics reported in this study are for a model trained on seven cancer subtypes and a single exome capture kit and sequencing center. A model trained on more diverse input data should generalize better than ours, and we expect our reported performance to increase with training set size and diversity. Training on a dataset with tumors of different subtypes, purity, ploidy, copy-number profiles, and mutational spectra, as well as multiple WES data sources will likely improve performance of a model applied to a cohort of contrasting biology and technical data quality. Conversely, analyzing a homogenous cohort (e.g., from a single cancer subtype) might benefit from training on a similar cohort, especially if including features like nucleotide substitution. In cohorts with a mix of matched-normal and tumor-only samples, it is possible to estimate the performance on the tumor-only subset in a way akin to the methodology outlined in this work, by running parallel matched-normal and tumor-only variant-calling pipelines on the matched-normal subset and evaluating the resultant classifications with the matched-normal truth labels.

Because the TabNet architecture was designed for tabular data, TabNet could be applied to a wide range of other tasks in genomics. It has been shown to outperform XGBoost in multiple classification tasks(Arik and Pfister 2019) and can be used for regression and multi-label classification. As we have demonstrated, TabNet’s feature masks enable interpretation of classification results. This is not possible with non-attentive deep learning methods(Chaudhari et al. 2019). The extent to which attentive models offer faithful explanation of predictions has been debated, working in some contexts and not others(Jain and Wallace 2019; Wiegreffe and Pinter 2019; Serrano and Smith 2019), but in this work, we see clear concordance between the values in the feature masks and statistical differences in the data.

For somatic mutation calling with DNA sequencing data, we expect no algorithm will ever be as good as having the matched normal. One can imagine a perfectly clonal, diploid tumor without copy number alterations and without any contaminating normal tissue (100% purity). In this hypothetical tumor, the variant allele fraction of somatic mutations would be distributed identically to the germline variants. Somatic mutations may exhibit characteristic genomic distribution and nucleotide substitution patterns(Alexandrov and Stratton 2014; Milholland et al. 2017), offering modest advantage for somatic vs germline classification in some tumor-types. Yet, without a substantial fraction of normal stromal tissue in the bulk WES biopsy, we expect methods based partly on VAF statistics such as ours will never perform as well as having the matched normal sample.

A major limitation in human genomics and precision medicine is that not all subpopulations are well-represented in genomic studies(Popejoy and Fullerton 2016). Human germline variant databases predominantly consist of subjects of White European ancestry, and this bias diminishes the reliability of naive tumor-only variant calling methods for Blacks relative to Whites(Asmann et al. 2021; Halperin et al. 2017). By integrating multiple informative features such as COSMIC and germline databases, variant allele fractions, and the local copy number ratios of known heterozygous germline SNPs, we can reduce much of this racial bias. We suspect this benefit conferred by our approach will improve further as germline variants from racial minorities become more represented in large genomic databases.

## Materials and Methods

### TCGA genomic data acquisition

Manifest files for downloading TCGA genomics data were generated using the TCGA-Biolinks(Colaprico et al. 2016) R package. 15 Patients for each indication were selected from TCGA randomly, conditioned on the patient having a single tumor-sample and single normal sample. 15 acute myeloid leukemia samples were originally included in the training set but were removed due to the presence of somatic mutations in the normal sample. BAMs were downloaded from GDC using the GDC Data Transfer tool. The samtools “collate” command was used prior to extracting FASTQs from the GDC BAMs.

### Hugo et al. metastatic melanoma WES data acquisition

Sequencing data from 23 metastatic melanoma patients sequenced at UCLA were downloaded from SRA using SRA toolkit and the command fastq-dump -split-3 –gzip $SRR. The 23-sample subset was chosen because it had available capture kit metadata.

### Alignment

FASTQs were aligned to hg38 using the Sentieon implementation of BWA-MEM(Freed et al. 2017). We used a consistent bioinformatics approach across all batches and cohorts.

### Panel of normals construction

**A** panel of normals is routinely used in whole exome sequencing analysis to filter out germline SNPs and alignment and technical artifacts inherent to the capture-kit choice. It is also used for CNV analysis – the germline copy number of many samples are used to capture-kit-specific depth biases. A leave-one-out panel of normals strategy was chosen to maximize the number of normal samples available for training. A further benefit of the method is it ensures that the racial demographics of the normal samples in the panel are representative of the cohorts used in training and evaluation.

A separate leave-one-out panel of normals was constructed for each of the 195 TCGA patients in this study. For a given capture kit with N patients sequenced, the leave-one-out approach is as follows: for each of the N patients, gather the N -1 normal samples from every other patient, and use these N - 1 normal samples to create both the CNV log2-copy number reference and normal panel VCF (VCF panel of normals). This strategy is analogous to leave-one-out cross-validation. The CNV and VCF panel of normals from TCGA data were matched with the capture kit of the tumor samples. For the 23 metastatic melanoma samples, CNV and VCF normal panels were both derived from a randomly chosen patient from the Nimblegen SeqCap EZ Exome v3 sequenced TCGA cohort.

### Variant level panel of normals construction

BCFtools(Danecek et al. 2021) and the merge command was used to aggregate the germline VCFs of the N-1 normal samples. All identified variants occurring in at least two of the samples were added to the normal panel VCF.

### Copy Number panel of normals construction

CNVkit(Talevich et al. 2016) output .cns files that were aggregated using the command. The bins were specified using the capture kit’s baits BED file, lifted over from hg19 to hg38 with the UCSC LiftOver tool.

### Copy number calling

We used CNVkit to generate log2 copy number ratios and segments using the circular binary segmentation algorithm. Batch mode for single TCGA samples the leave-one-out panel of normal were used. For the Hugo melanoma cohort, batch mode was also used, with the CNV panel of normals from a randomly chosen TCGA patient.

cnvkit.py batch $tumor -r $pon -p $procs_per_job --output-dir $sample

cnvkit.py call $sample/$sample\_tumor.cns -o $sample/$sample.call.cns

### Variant Calling

Sentieon’s TNScope(Freed, Pan, and Aldana 2018) was applied to the hg38-aligned BAMs and the capture-kit-matched panel of normals. No patient-matched normals were included in the process-matched panel of normals. SnpSift v4.3 added dbSNP(Sherry et al. 2001) build 151 and COSMIC(Tate et al. 2019) v85 annotations to all VCFs with the following command: SnpSift Annotate -a $COSMIC_VCF $SNP_EFF_ANNOTATED_VCF

dbNSFP4.0(Liu et al. 2020) was used to annotate variants with databases such as 1000 Genomes(1000 Genomes Project Consortium et al. 2015) and ExAC(Lek et al. 2016). We constructed ‘pop_max’, a single aggregate feature derived from dbNSFP for filtering and the machine learning model. pop_max, calculated by taking the maximum population allele frequency across the following dbNSFP databases: 1000Gp3_AF, TWINSUK_AF, ALSPAC_AF, UK10K_AF,ExAC_AC,ExAC_AF,gnomAD_exomes_AF, gnomAD_genomes_AF.

### Variant filtering

A set of criteria was chosen for pre-filtering variants such that artifacts and common germline SNPs are eliminated before applying training or applying the tumor-only classifier. These eliminated variants do not count as true negatives, thus our specificity and NPV are calculated conservatively. The criteria isolated passing, coding mutations for all tumor-only variant calls and is as follows: population allele frequency < 0.01 across the 8 germline databases, SnpEff annotation ontology in missense, nonsense, frameshift_indel, inframe_indel, FPfilter == ‘PASS’, Sentieon TNScope filter == ‘PASS’.

FPfilter(Koboldt et al. 2012) eliminated sequencing and alignment artifacts. TNScope filter flags likely sequencing errors (using the t_lod_fstar of Mutect 2) as well as artifacts and germline mutations identified with the process-matched panel of normals. We discarded these variants and kept only the variants that we’d consider to be somatic coding mutations.

### PureCN

We ran PureCN using the production configuration recommended in the official documentation. For input we used the COSMIC and dbSNP-annotated tumor-only VCFs after removing artifacts from the VCFs using bcftools (TNScope filter == ‘PASS’). used with PureCN. normalDB was constructed for every PoN VCF used in this study with the command Rscript $PURECN /NormalDB.R --outdir $out_dir --normal_panel $pon_vcf --assay $patient_id + --genome hg38 --force

The copy-number ratio .cnr files from CNVkit were converted to segmentation files (.seg) using the ‘cnvkit export’ command. The hg38_simple_repeats.bed file was downloaded from UCSC to blacklist SNPs in tandem repeat regions(Benson 1999). 250 cores were used per sample and the “—postoptimize” flag was turned on. The full command is as follows:

Rscript $PURECN --version; Rscript $PURECN --out $out_dir --sampleid $patient_id –tumor $COPY_NUMBER_RATIO --segfile $seg_file --mappingbiasfile $normal_db --vcf $vcf -- snpblacklist $simple_repeats --genome hg38 --parallel --cores 250 --funsegmentation Hclust -- force --postoptimize --seed 123

### Comparing TabNet and PureCN

Unfiltered variants from our variant calling pipeline were merged with the classified variants from PureCN. Variants were subsequently filtered using the same criteria and thresholds that we applied to isolate coding somatic mutations, including the TNScope filter, FPFilter, coding mutation ontology, and population database frequency. TabNet predictions were merged, and call rate was assessed for TabNet and PureCN by calculating the number of variants with posterior somatic probability predictions. True positives were defined as: somatic mutations correctly classified as somatic, false positives: rare germline variants misclassified as somatic, false negatives: true somatic mutations misclassified as germline, true negatives: rare germline variants correctly classified as germline.

### TabNet training

The open-source repository PyTorch TabNet (https://github.com/dreamquark-ai/tabnet) was adapted for, with PyTorch version 1.7.0.

The following model hyperparameters were used to build a TabNet network: n_d = 24, n_a = 24, n_steps = 4, gamma = 1.5, n_independent 2, n_shared = 2, lambda_sparse = 0.0001, momentum = 0.3, clip_value = 2.

Training was achieved for 100 epochs, using the Adam Optimizer with a learning rate of 0.02, and a batch size of 4000 and virtual batch size of 256. Although TabNet does not require categorical features to be one-hot-encoded, we did this to allow for more flexibility with other machine learning models, such as a random forest. A custom loss function was designed to maximize the average precision score (the area under the precision recall curve). We trained the model for a total of 100 epochs, after which we selected the model from the epoch with the best performance on the validation set. We repeated the train-validate-test process three times to ensure reproducibility of this training strategy. Training completed in less than 2 hours on the CPU, and approximately 15 minutes using GPU acceleration with NVIDIA P100. GPUs were not used for the time benchmark comparison for TabNet and PureCN.

### Regression and covariance analysis

Linear regression and covariance calculations were calculated using the R computing environment, version 3.5.2. To calculate quartiles in Supplemental Figure 4A., we used bootstrap sampling with 1000 bootstrap replicates.

### TMB calculation

There is no consensus on how to normalize TMB, i.e., whether to use the size of the exome in the human genome or the size of the regions targeted by the exon capture kit. Often the TMB is presented without normalizing. We obtained exon target .BED files for the three capture kits in this study from the manufacturer websites. We used the UCSC liftOver tool to convert them from hg19 to hg38. The total footprint of the exon targets from SeqCap EZ HGSC VCRome, and Nimblegen SeqCap EZ Exome v3 kits, was 33.0, 37.3, and 63.5 megabases, respectively. Since we used three distinct exon capture kits in this study, for simplicity we decided to normalize the total somatic mutation count across all datasets by dividing by a constant factor: 41, corresponding to the patient-weighted average of the three kits’ target footprint size in megabases.

## Code Availability

This tumor-only somatic-germline classifier was written in Python. All code including feature engineering and model training and evaluation is available at https://github.com/AbbVie-GRC-Methods-Dev/new_normal and is usable under the MIT license.

