## Supplemental Materials for "Attentive deep learning-based tumor-only somatic mutation classifier achieves high accuracy agnostic of tissue type and capture kit"

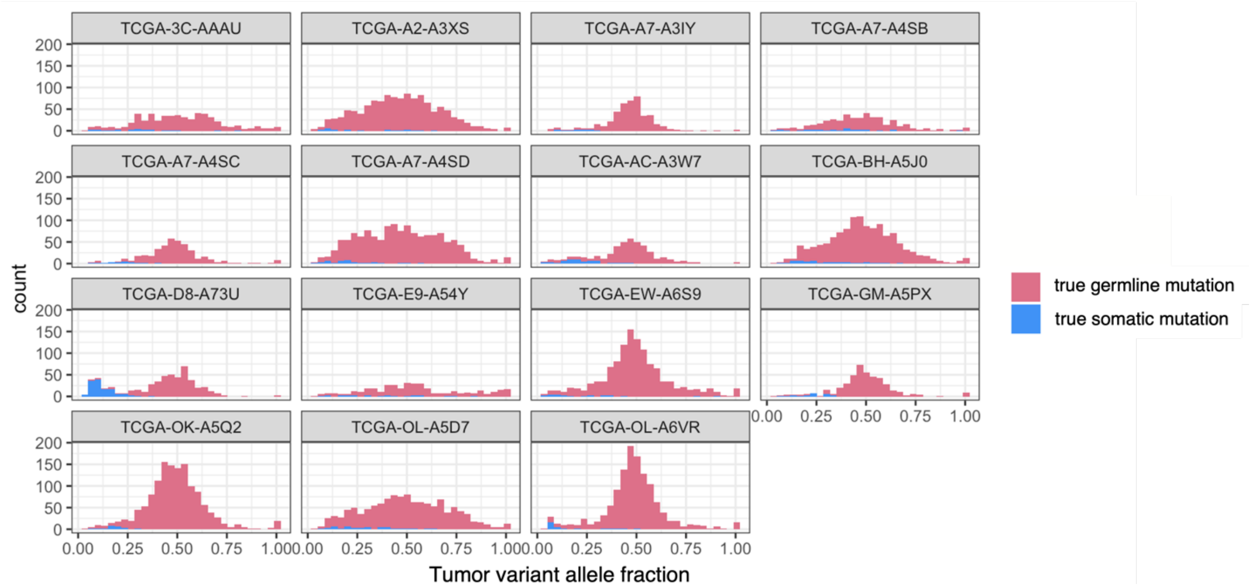


**Supplemental Figure 1:** The overlap between distributions of variant allele fractions (VAFs) of true somatic and true rare germline variants is a source of complication in tumor-only somatic variant calling. Rare germline variants greatly outnumber true somatic mutations, leading to a high false positive rate in tumor-only somatic variant calling, Data shown are for the 15 breast cancer patients from TCGA included in this study.

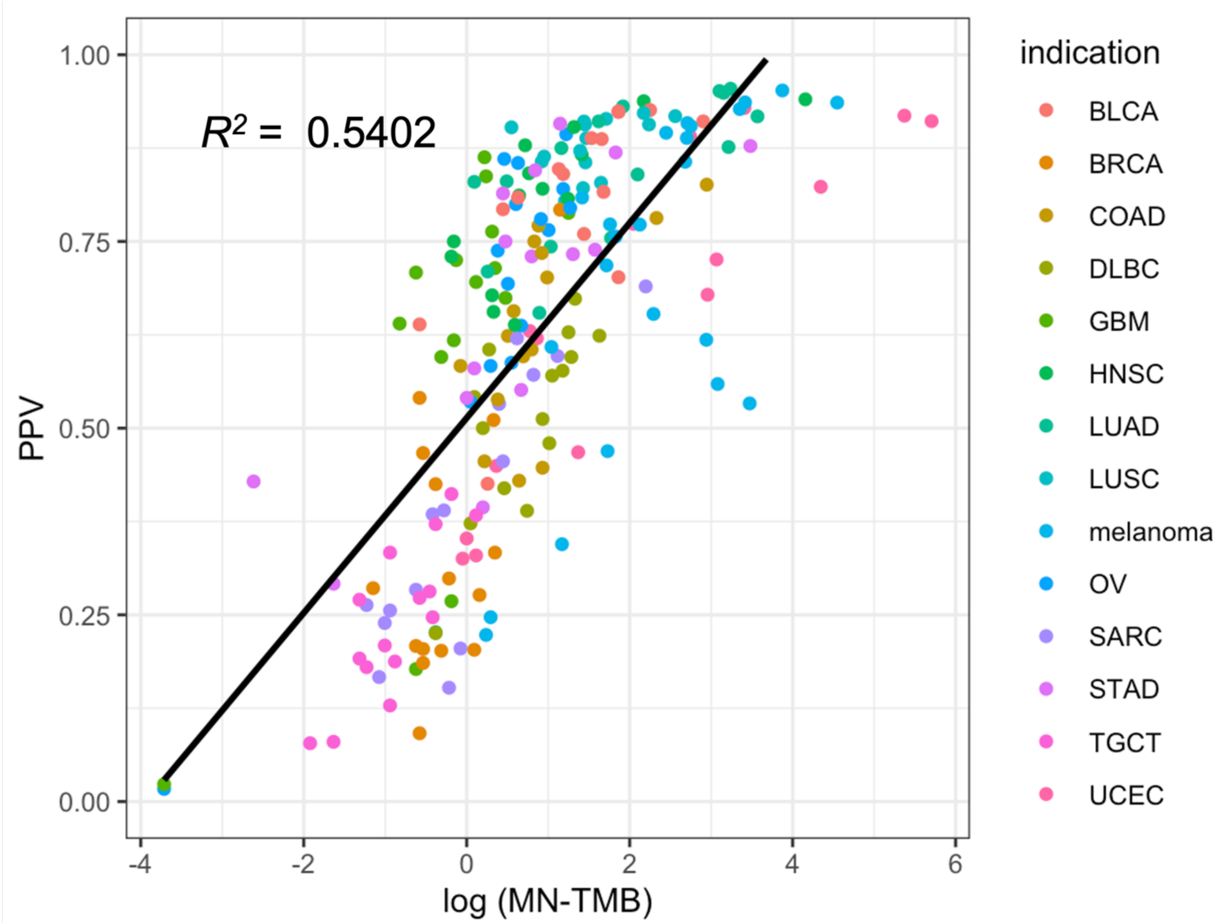


**Supplemental Figure 2:** Across samples, the positive predictive value (PPV) of the somatic-germline classifier is correlated with the log matched-normal TMB (MN-TMB, the "true" TMB). All 218 samples in this study are displayed.

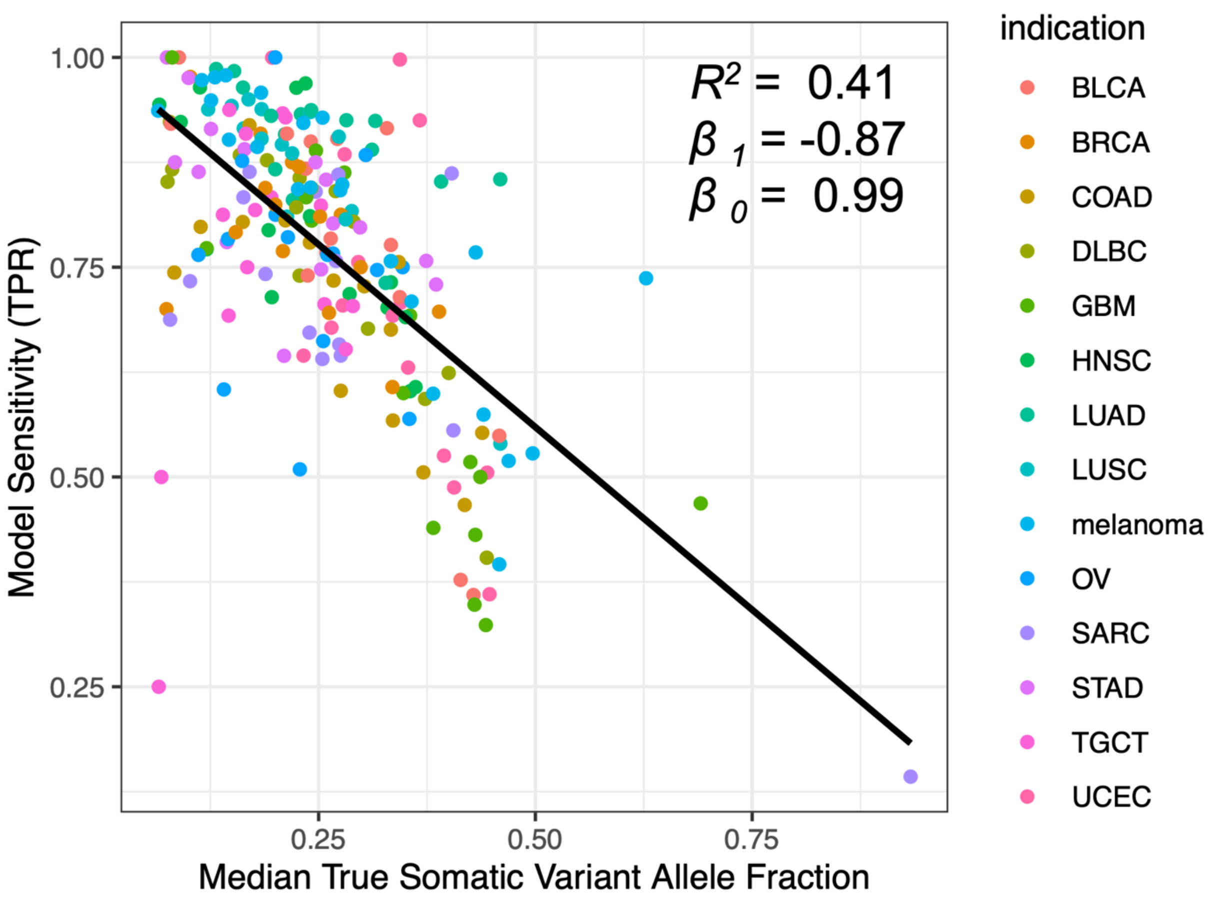


**Supplemental Figure 3:** Classifier **s**ensitivity (TPR) is negatively correlated with the median variant allele fraction (VAF) of the true somatic mutations (MVTSM), a metric related to tumor purity. All 218 samples in this study are displayed.

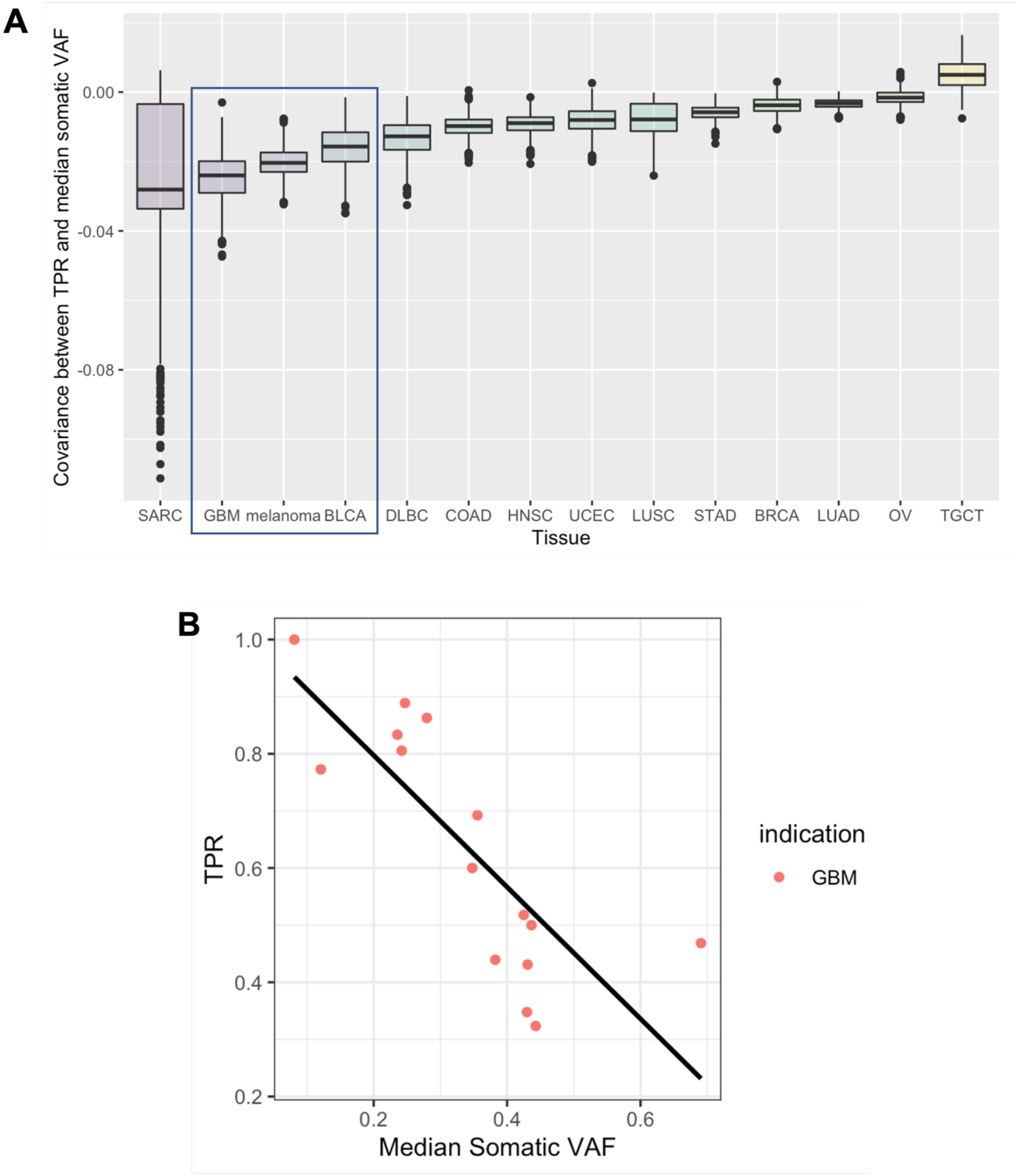


**Supplemental Figure 4: A)** Bootstrapped covariance analysis ranks the 14 cancer indications in this study by the strength of the inverse relationship between sensitivity (TPR) vs median VAF of true somatic mutations (MVTSM). **B)** Glioblastoma multiforme (GBM) is highlighted as a cancer subtype whose TPR is strongly inversely related to MVTSM.
